# Efficient single copy integration via homology-directed repair (scHDR) by 5’modification of large DNA donor fragments in mice

**DOI:** 10.1101/2021.09.30.462539

**Authors:** Rebekka Medert, Thomas Thumberger, Tinatini Tavhelidse, Tobias Hub, Tanja Kellner, Yoko Oguchi, Sascha Dlugosz, Frank Zimmermann, Joachim Wittbrodt, Marc Freichel

## Abstract

CRISPR/Cas approaches have largely replaced conventional gene targeting strategies. However, homology-directed repair (HDR) in the mouse genome is not very efficient, and precisely inserting longer sequences using HDR remains challenging, given that donor constructs preferentially integrate as concatemers. Here, we show that injecting 5’biotinylated donor DNA in mouse embryos at the two-cell stage leads to efficient single-copy HDR (scHDR) alleles. Our dedicated genotyping strategy showed that these alleles occurred with a frequency of 19%, 20%, and 26%, respectively, in three independent gene loci, indicating that scHDR is dramatically boosted by 5’biotinylation. Thus, we suggest that a combination of a 5’biotinylated donor and diligent analysis of concatemer integration are prerequisites for efficiently and reliably generating conditional alleles or other large fragment knock-ins into the mouse genome.

## INTRODUCTION

In the past, gene targeting approaches in embryonic stem cells (ESC) have been used to integrate foreign DNA fragments at specific gene loci in the mouse genome (1). This approaches were used to generate mouse lines expressing proteins with e.g. fluorescent tags or to introduce loxP-flanked exons to allow gene deletion at specific time points and in specific cell types. Exogenous DNA was integrated into the mouse genome via homologous recombination (HR) (2–4). Although the ESC method revolutionized genome engineering, the approach requires screening to correctly target ESCs, and this screening is time and labor intensive. However, screening is necessary because HR efficiency is often under 1% (1,5). Genome editing approaches in mammalian cells, including mouse embryos, allow the deletion and introduction of point mutations in the mouse genome via CRISPR/Cas9-mediated techniques. This approach is highly efficient and has replaced classical ESC approaches in recent years. However, introducing double strand breaks (DSBs) increases the frequency at which homologous DNA fragments integrate at a given locus in the genomes of eukaryotes (6,7). The integration of larger (>100bp) homologous donor fragments at precise genomic locations in the mouse genome via homology-directed repair (HDR) following co-injection of *Cas9* mRNA, sgRNAs, and DNA vectors was reported early in the development of this technology (8). However, such events are fairly infrequent since most DSB events are still repaired using error-prone non-homologous end joining (NHEJ) mechanisms rather than by HDR. NHEJ can result in insertions and deletions (INDELS). In addition to HDR-mediated repair, several alternative approaches have been described to knock-in DNA fragments, such as homology independent targeted integration (HITI) (9) and microhomology-mediated end joining (MMEJ) (10). Approaches that can facilitate the knock-in of DNA fragments into the mouse genome after CRISPR/Cas9 induced DNA DSB include CRIS-PITCh (11), Tild-CRISPR (12), Easi-CRISPR (13), 2C-HR-CRISPR (14), and CRISPR-READI (15). These methods all use different donor strategies and different DNA repair mechanisms (16). For example, CRIS-PITCh uses a donor plasmid that is linearized in vivo by Cas9. Short homology arms (20-40 bp) are used to enable knock-in via microhomology mediated end-joining (MMEJ). The knock-in efficiency of a CRIS-PITCh-generated floxed mouse line described by Aida et al. is around 35.7% (11). In an alternative method, Easi-CRISPR, long single-stranded DNA donors are used to knock-in reporters or recombinases and loxP-flanked fragments via HDR. The knock-in efficiency of the transgene has been reported to be as high as 30% to 60%; it can even reach 100% for some loci (17).

Linear dsDNA injected into mammalian cells multimerizes via head-to-tail joining and may integrate into the genome as a concatemer (2,18,19). In previously published knock-in studies, little attention has been paid to the formation of donor multimers, which can integrate at the target site or at other random sites in the mouse genome. A recent study showed that high background rates of multimer insertions of donor DNA confound the generation of properly targeted alleles (20). Such events may occur due to numerous mechanisms and combinations of HDR and NHEJ; in a generation of six different conditional knockout lines reported by Skryabin et al., 57% of the F1 mice were positive for multiple copy head-to-tail integrations (20).

We show that this feature of dsDNA multimerization is not exclusive to mammalian cells; it also occurs when linear dsDNA is injected into zygotes of the Japanese rice fish medaka (*Oryzias latipes*). This suggests that there is a general mechanism for processing linear dsDNA upon delivery into cells. We also found that certain modifications to the 5’ end of DNA strands protect them from multimerization and lead to a significantly higher single-copy HDR (scHDR) efficiency compared to unmodified DNA ends (21). We worked to balance protection from donor concatemerization and single-copy HDR efficiency against embryo survival rates and achieved the best results from donor modification by using 5’biotinylation on the ends of the dsDNA donor. However, DSB repair mechanisms differ substantially across species. Therefore, it remained to be shown whether a 5’modification to the doner dsDNA also improves HDR efficiency in mice. The present study evaluated this in mouse embryos.

Interestingly, Gu et al. showed that a 5’biotinylated dsDNA PCR donor combined with a Cas9-Streptavidin fusion protein (Cas9-mSA) increased HDR efficiency by up to four times in mouse embryos compared to an unmodified dsDNA PCR donor (14). That study attributes this increased HDR efficiency to the interaction of the donor-Biotin Cas9-Streptavidin, which results in more efficient recruitment of the biotinylated donor to the Streptavidin-linked Cas9 protein. However, that study does not address whether this increased HDR efficiency occurs by prevention of donor multimerization, which has proven to be efficient in medaka, independent of Streptavidin coupling to the Cas9 endonuclease (21).

The present study confirms the formation of highly efficient single-copy HDR (scHDR) upon injection of 5’biotinylated dsDNA donor into mouse embryos at the two-cell stage. This was done to establish conditional mouse lines at three different loci; loxP-flanked alleles of the *Akr1b3*, *Fam129b*, and *Jpt2* genes were generated. To address pitfalls such as multimer integration, we developed a tandem PCR-based genotyping assay. Our study reveals that an approach that combines the use of 5’biotinylated donor DNA and careful analysis of concatemer integration is an efficient, reliable strategy for obtaining mice with precise large gene modifications (e.g., conditional alleles).

## RESULTS

### Strategy to achieve and validate CRISPR/Cas9 mediated single-copy homology-directed repair (scHDR) of long donor fragments in the mouse genome using 5’biotinylated donor sequences

We set out to generate conditional alleles to establish mouse models that allow time-dependent and cell type-specific gene deletions. To this end, we aimed to use a donor DNA template containing a loxP-flanked target exon that integrates at the target locus after CRISPR/Cas9 induced DNA double-strand breaks (DBS) with sgRNAs cutting in the introns 5’ (sgRNA #1) and 3’ (sgRNA #2) of the target exon, respectively (Figure 1A). HDR-mediated donor insertion should be facilitated using 5’ and 3’ homology arms of about 500 bp (Figure 1A). However, the targeted integration of a donor using homology-directed repair can frequently lead to the insertion of multimers (20,21). In fish, the multimerization of donor DNA can be efficiently reduced by 5’ modification, which favors scHDR (21). To determine whether this also applies to a mammalian system, we 5’biotinylated DNA donors for scHDR in mouse embryos (Figure 1B).

**Figure 1.**
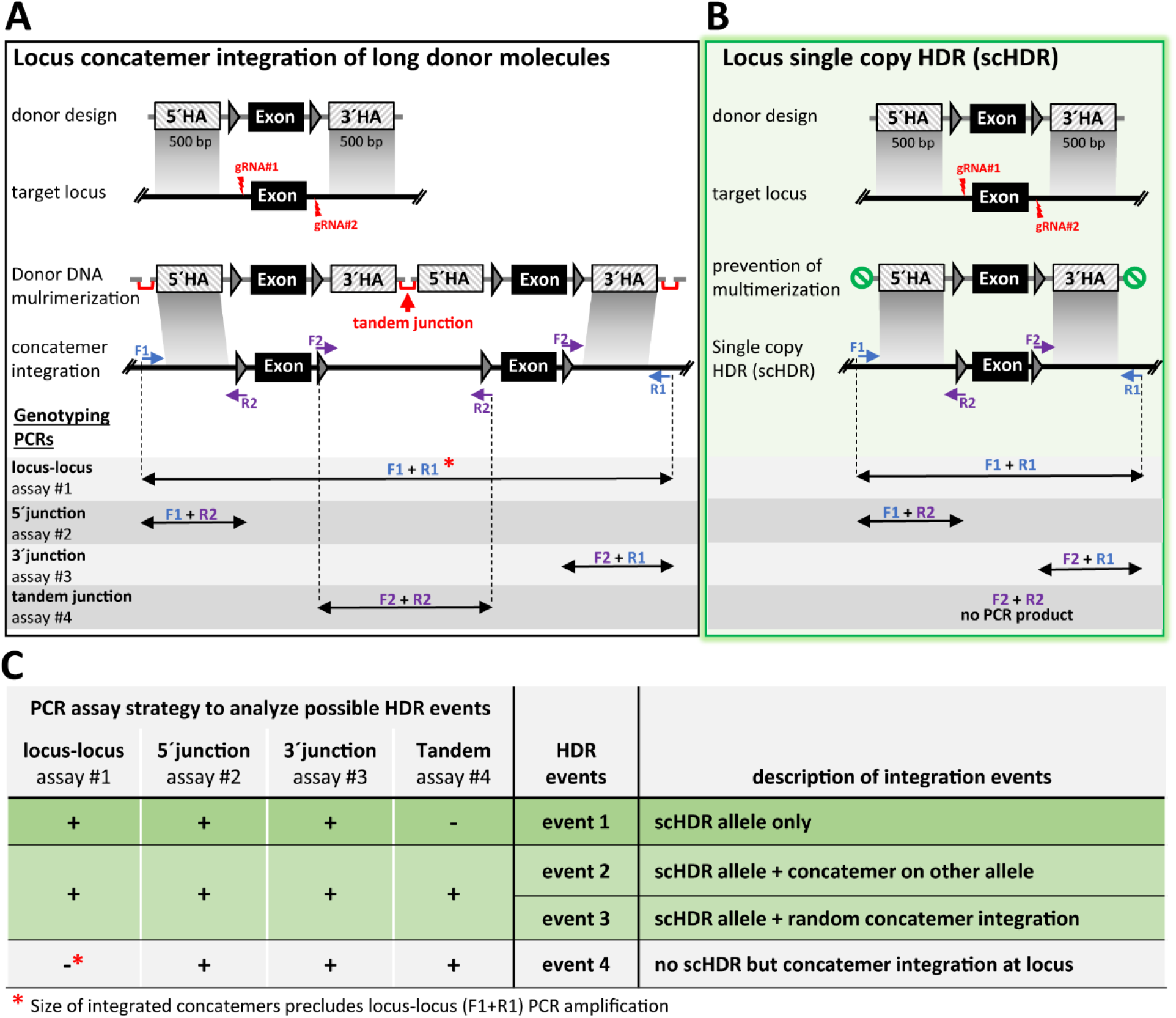
CRISPR/Cas9-mediated two-sgRNA HDR strategy to generate conditional alleles. **(A-B)** Targeting strategy for integration of a PCR generated dsDNA donor containing the target exon flanked by loxP sites (triangles) and 500 bp homology arms (HA). **(A)** unmodified PCR donor molecules multimerize and integrate as concatemers at the locus of interest. **(B)** 5′biotinylation of PCR donor prevents donor multimerization and allows precise single-copy HDR (scHDR) of the donor template. Genotyping PCR assays identifying precise scHDR (locus-locus PCR, assay #1), targeted integration (5′junction PCR, assay #2 and 3′junction PCR, assay #3) and concatemer integration (tandem junction PCR, assay #4) of the donor template. Tandem junction PCR strategy was used to identify the integration of donor concatemers in the mouse genome. The forward primer (F2) is located at the 3′end of the donor, whereas the reverse primer (R2) is located at the 5′end. Donor concatemer integration results in a PCR product whereas correct scHDR results in no tandem junction PCR product. **(C)** PCR assay strategies comprised of PCR assays #1-#4 to detect scHDR alleles with and without random or targeted concatemer integration. Application of this diligent analysis allows identification of defined integration events of the donor template in the mouse genome.

Since such unintended integration events cannot be detected by PCR assays that are restricted to the 5’ and 3’ junctions of the integration site, we implemented a genotyping protocol consisting of four PCR assays (Figure 1A). This protocol enabled us to distinguish the desired single-copy donor integration via HDR (scHDR) from other integration events. PCR assay #1 was a locus-locus PCR (Figure 1A, blue arrows); the primers were placed outside the homology arms, making it possible to discriminate alleles that underwent scHDR from unedited wild type alleles, alleles with short indels, and alleles arising from the deletion of DNA sequences between sgRNA target sites. These deletions are caused by the introduction of a DSB followed by a NHEJ-mediated repair without integration of the donor. PCR assay #1 also enabled us to identify concatemers of donor DNA. However, larger fragments derived from sequential fusions of the donor fragment (dimers or larger multimers) are less likely to be amplified by the locus-locus PCR because of its decreased amplification efficiency for larger template sizes. The assay’s efficiency could also be negatively impacted by the presence of competing shorter DNA amplicons arising from wild type and/or knockout (deletion of exon) alleles. A 5’ junction PCR (assay #2) and a 3’ junction PCR (assay #3) are often used for genotyping and can help confirm the targeted integration at the locus. However, these assays cannot rule out donor concatemer integration since the integration of multimers can also take place through HDR. Therefore, 3’ and 5’ junction PCR products of the same size arise (Figure 1C).

A tandem junction PCR (assay #4) was used to identify the multimeric integration events of donor fragments. The forward (F2) and reverse (R2) primers which face in opposite directions (Figure1A, purple arrows) were used for this assay. The possible integration events and the expected results of the tandem junction PCR assay are summarized in the table shown in Figure 1C and Supplementary Figure 1. To summarize, the lack of additional unwanted integration in the presence of scHDR (event 1) is revealed by the presence of amplification products of the expected size in PCR assay #1 (locus-locus), PCR assay #2 (5’ junction), PCR assay #3 (3’ junction) and by the absence of an amplicon in PCR assay #4 (tandem junction) (Figure 1C, upper row). If a PCR product is obtained in assay #4 (tandem junction) and the results of assays #1 to #3 are positive, this indicates multimer integration arising from scHDR integration combined with either random integration (event 2) or scHDR integration in one allele along with concatemer integration at the target locus of another allele (event 3). A positive tandem junction PCR (assay #4) and an absence of amplicons in the locus-locus PCR (assay #1) indicates concatemer integration at the target locus if the 5’ and 3’ junction PCRs (assays #2 and #3) are positive (event 4). A positive assay #4 combined with a lack of amplicons in assay #1 indicates random integration if the 5’ and 3’ junction PCRs (assays #2 and #3) are negative. Possible variants of concatemer integration events are illustrated in Supplementary Figure 1, but even more incorrect integration events can result from the degradation of the locus and/or donor sequences and from various combinations of HDR and NHEJ (20).

### 5’ biotinylation of DNA ends ensures the monomeric state of donor molecules *in vivo*

We evaluated the multimerization of PCR donor molecules in medaka and mice *in vivo*. Unmodified donor molecules (Akr1b3-flox, Supplementary Figure 3) and donors modified with 5’ biotinylation were microinjected into medaka zygotes (at the one-cell stage) and mouse embryos (at the two-cell stage) (Figure 2A, B). One and six hours after the microinjection, donor multimer formation was evaluated using droplet digital PCR (ddPCR) to identify tandem junctions (Figure 2C, probe 2 assay, which yielded FAM-positive droplets) resulting from the ligation of at least two donor molecules. The number of tandem junctions was normalized to the total number of donor molecules (assessed with a probe 1 assay, which yielded HEX-positive droplets). We ruled out the possibility that the donor molecules had already formed multimers outside the cytosol, as no amplicons were obtained from the tandem junction (probe 2) ddPCR assay in not injected control medaka or mouse embryos. This was measured by analyzing samples derived from unmodified donor molecules released outside the embryos (Figure 2D-G, left panels). In stark contrast, high numbers of donor multimers were detected one hour after the injection of unmodified donor molecules. In medaka embryos, this number was decreased one hour and six hours after the injection of 5’biotinylated donor molecules (Figure 2D, E). In mouse embryos, multimer formation was almost entirely prevented at both time points by the 5’biotinylation of donor molecules (Figure 2F, G).

**Figure 2.**
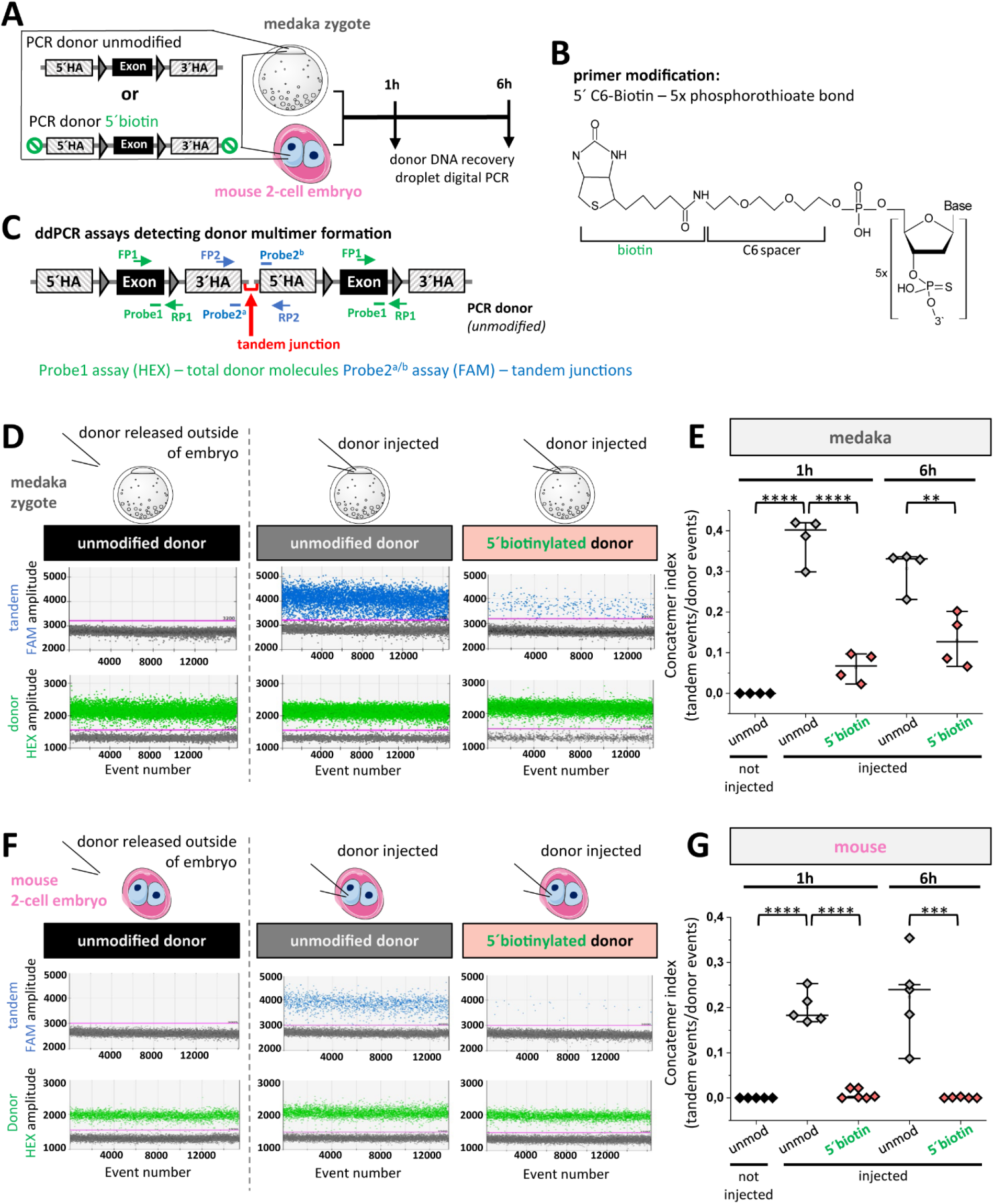
Biotinylation of DNA 5′ends ensures monomeric state of dsDNA donor molecules in medaka and mouse embryos. **(A)** Unmodified versus 5′ biotinylated PCR fragments were injected in medaka or mouse embryos and the donor multimerization state was analyzed one and six hours post injection. **(B)** Structure of 5′biotinylated primer modification for PCR donor generation. **(C)** Droplet digital PCR (ddPCR) assays to quantify total donor molecules (Probe1 assay, HEX) and tandem junctions (Probe 2 assay, FAM) resulting from donor multimerization. **(D)** Multimerization state of not injected unmodified donor molecules (black) compared to unmodified (grey) and 5′biotinylated (bright red) donor molecules one hour after injection into medaka embryos. Droplet count view shows total donor molecule (green) and tandem junction (blue) event numbers. **(E)** Quantification of multimerization state of unmodified and 5′biotinylated donor molecules in medaka embryos one and six hours post injection. **(F)** Multimerization state of not injected unmodified donor molecules (black) compared to unmodified (grey) and 5′biotinylated (bright red) donor molecules one hour after injection into mouse embryos. **(G)** Quantification of multimerization state of unmodified and 5′biotinylated donor molecules in mouse embryos one and six hours post injection. Concatemer index: tandem junction event number/donor molecule event number. P values were calculated by two-sided t-test, *P < 0.05, **P < 0.01, ***P < 0.001, ****P < 0.0001; n = 4-5 biologically independent replicates. Error bars represent mean ± SD. Mouse embryos were drawn by using image elements from Servier Medical Art. Servier Medical Art by Servier is licensed under a Creative Commons Attribution 3.0 Unported License (https://creativecommons.org/licenses/by/3.0/)

### 5’biotinylation of donor DNA facilitates scHDR

Next, we investigated whether 5’biotinylation of the PCR donor allowed efficient scHDR integration in the mouse genome. First, we aimed to generate a conditional *Akr1b3* allele to establish a mouse model in which *Akr1b3* could be deleted in a time-dependent and cell type-specific manner. To this end, exon 3 needed to be flanked by loxP sites using two sgRNAs that cut in the introns 5’ (sgRNA #1) and 3’ (sgRNA #2) of exon 3, respectively (Figure 3A). To insert loxP sites, the endogenous sequences encoding exon 3 should be replaced by a 1.9 kb PCR donor fragment containing a loxP-flanked exon 3, a 400 bp 5’ homology arm, and a 500 bp 3’ homology arm (Figure 3A). To boost HDR-mediated insertion, the injections were performed at the two-cell stage with a long G2 phase (14). In the first set of experiments, the embryos were injected with a CRISPR mix containing *Cas9* mRNA, sgRNAs, and 5’biotinylated donor DNA (Supplementary Figure 3A). The control included three groups: non-injected embryos (Supplementary Figure 3B, lane 1), embryos injected with pure injection buffer (Supplementary Figure 3B, lane 2), and embryos injected with a mix that did not contain any donor DNA (Supplementary Figure 3B, lane 3). Five of fifteen independent pools of E4.5 embryos injected with the 5’biotinylated donor were positive for scHDR in the *Akr1b3* locus, demonstrated by a 1.8kb amplicon in the locus-locus PCR assay #1 (scHDR efficiency 33%, Supplementary Figure 3B, lane 4 and Supplementary Figure 3C).

**Figure 3.**
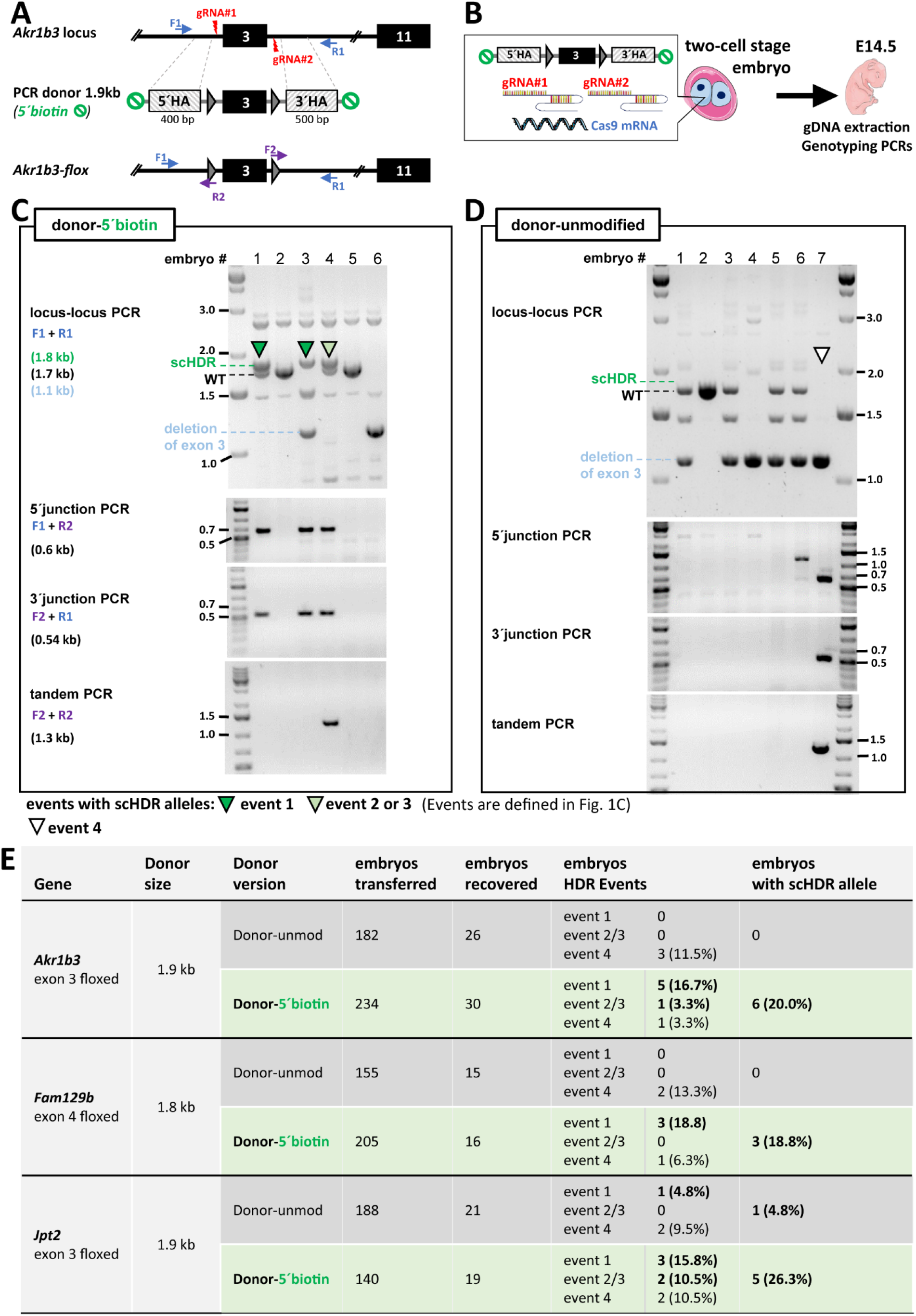
Donor 5′biotinylation achieves efficient single-copy HDR (scHDR) in mouse. **(A)** Two-sgRNA targeting strategy for scHDR of a PCR donor template containing exon 3 flanked by loxP sites. **(B)** two-cell stage microinjection strategy to deliver *Cas9* mRNA, sgRNAs and PCR donor-5‘biotin or donor-unmodified into the mouse embryo followed by genotyping of single embryos at embryonic stage E14.5. **(C)** CRISPR/Cas9 mediated scHDR at the *Akr1b3* locus is only achieved when PCR donor-5’biotin is injected. **(D)** Genotyping PCRs identified tandem junctions but no scHDR events in the presence of PCR donor-unmodified. **(E)** Table showing the occurrence of HDR events 1-4 and scHDR efficiency for PCR donor-5‘biotin and donor-unmod at different loci. scHDR efficiency was evaluated at embryonic stage E14.5. Nucleotides and mouse embryos were drawn by using image elements from Servier Medical Art. Servier Medical Art by Servier is licensed under a Creative Commons Attribution 3.0 Unported License (https://creativecommons.org/licenses/by/3.0/).

To determine whether the high scHDR efficiency was due to the 5’modification of the PCR donor, we compared injections of modified and unmodified PCR donors (three different injection sessions). To analyze each injected embryo individually, the embryos were transferred to a foster mouse and isolated on embryonic day E14.5 (Figure 3B). Locus-locus PCR identified scHDR exclusively when PCR donor 5’biotin was injected (Figure 3C, dark green arrowheads, embryo #1 and #3). Although the detection of amplicons of the correct size (1.8 kb) in the locus-locus PCR (assay #1) indicates scHDR, the correct HDR-mediated integration of the donor can be accompanied by additional integrations of donor concatemers, indicated by a positive tandem junction PCR (assay #4). This was observed in embryo #4 (light green arrowhead, Figure 3C). A positive tandem assay can arise from additional integration at the second *Akr1b3* allele (event 2) or at random sites in the genome (event 3). The use of 5’biotinylated donor DNA resulted in scHDR alleles in 20.0% of the E14.5 embryos; this included embryos with event 1 (16.7%) and events 2 or 3 (3.3%) (Figure 3E). Embryos injected with the unmodified donor DNA showed no scHDR integration, although targeted multimer integration was identified via 5’ and 3’ junction PCRs and via tandem PCR (embryo #7 in Figure 3D).

To investigate whether 5’biotinylation of donor DNA allows efficient scHDR integration of donor molecules at other gene loci as well, we targeted the *Fam129b* locus, using a similar strategy to generate a floxed exon 4. We detected scHDR alleles in 18.8% of E14.5 embryos injected with a 5’biotinylated donor (size 1.9 kb); embryos injected with the unmodified donor showed only concatemeric integration (Figure 3E and Supplementary Figure 4A-C). A comparable targeting strategy using a 1.8 kb donor was then applied at a third gene locus, *Jpt2*. Here, scHDR alleles were observed at a frequency of 26.3% when the modified donor was used. In contrast to the other targeted loci, the unmodified donor injections also resulted in 4.8% scHDR at *Jpt2* (Figure 3E Supplementary Figure 4D-F). While the simple biotinylation of 5’ DNA donor ends dramatically boosts scHDR efficiency, scHDR only occurs occasionally and at low levels with unmodified donors; it is also locus dependent.

### Efficient germline transmission of alleles derived from scHDR

The 5’ biotinylated PCR donor was used to establish *Akr1b3^+/fx^* mouse lines. The CRISPR mix, which contained *Cas9* mRNA, sgRNAs, and the 5’biotinylated PCR donor, was again injected into embryos at the two-cell stage; the embryos were then transferred into foster mice (Figure 4A). PCR genotyping identified scHDR in six of the 28 F0 mice that were born (21%), and precise scHDR was confirmed by sequencing the locus-locus PCR product; this sequencing covered the entire fragment integrated into the *Akr1b3* locus, including the loxP sites (founder #39, Supplementary Figure 5). Tandem PCR showed that five of the six founders exhibited scHDR without any concatemer integration (scHDR only, event 1, dark green arrowheads, Figure 4B) and that only one of them exhibited additional concatemers (founder #23, light green arrowhead in Supplementary Figure 6) corresponding to integration event 2 or 3 (table, Figure 1C). scHDR-positive founders were crossed with C57BL/6N mice, and the F1 offspring were tested for germline transmission. PCR genotyping revealed germline transmission in the F1 generation in three founders; already after the analysis of two litters from each founder were analyzed (8/13 scHDR-positive offspring for founder #23; 7/17 for founder #38; 10/15 for founder #39, Figure 4C). Offspring with concatemer integration were only observed after founder #23 was mated, indicating integration event 3 (light green arrowhead with black stripes in Suppl Figure 6C). Nevertheless, founder #23 also generated numerous offspring presenting only scHDR integration (event 1, dark green arrowhead in Suppl Figure 6C).

**Figure 4.**
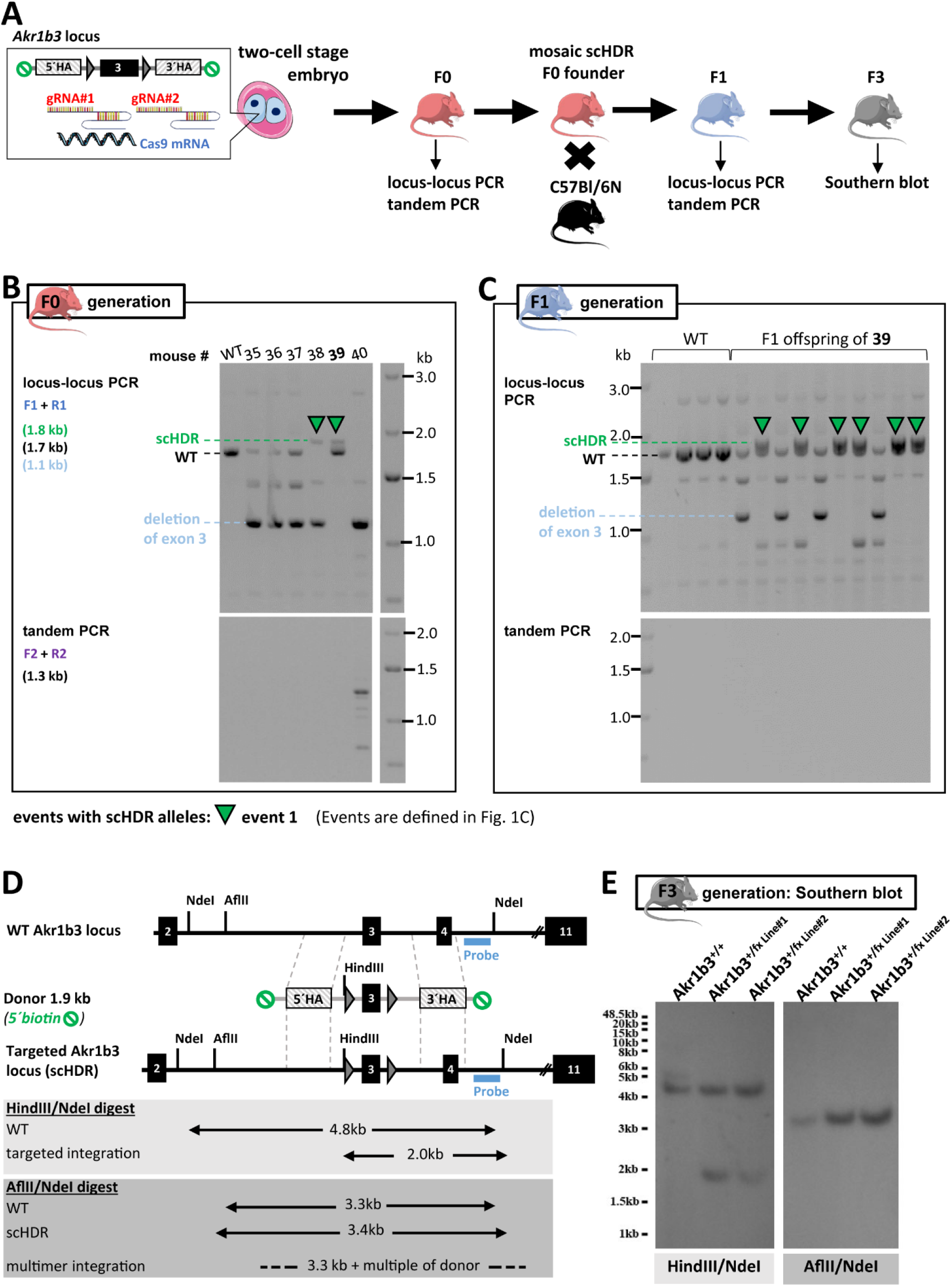
Efficient and stably transmitted *Akr1b3-flox* allele derived from scHDR. **(A)** two-cell stage microinjection strategy to deliver *Cas9* mRNA, sgRNAs and PCR donor-5‘biotin into the mouse embryo followed by genotyping of single founders using ear punch biopsies. Mating strategy of mosaic scHDR positive founder and genotyping of F1 generation. Southern Blot analysis in F3 generation confirms scHDR events. **(B)** Locus-locus PCR (assay #1) of F0 mice identified CRISPR/Cas9 mediated scHDR alleles (green arrowheads) upon injection of a PCR donor-5‘biotin. Tandem junction PCR of F0 mice identifies donor concatemer integration. **(C)** Locus-locus PCR of F1 generation identified germline transmission of scHDR alleles (dark green arrowheads). Tandem junction PCR excludes concatemer integration of the donor in scHDR positive F1 offspring. **(D)** Schematic representation of the murine *Akr1b3* gene targeting strategy indicating the location of the probe, as well as the restriction sites and the resulting DNA fragment sizes for **(E)** Southern Blot analysis of genomic DNA - left panel: heterozygous F3 Akr1b3^+/fx^ offspring shows the NdeI derived 4.8 kb wildtype band and demonstrates successful targeted integration of the donor by the specific band at 2.0 kb in case of the HindIII/NdeI digestion scheme. The Akr1b3^+/+^ littermate control lacks the donor encoded HindIII restriction site and shows only the NdeI derived 4.8 kb band. Right panel: cutting outside the donor cassette using AflII/NdeI results in a 3.3 kb Akr1b3 wildtype band and a 3.4 kb Akr1b3^+/fx^ band without any signs of larger concatemer bands at the *Akr1b3* locus. Nucleotides, mouse embryos and adult mice were drawn by using image elements from Servier Medical Art. Servier Medical Art by Servier is licensed under a Creative Commons Attribution 3.0 Unported License (https://creativecommons.org/licenses/by/3.0/)

These results highlight the need to carefully evaluate imprecise random or targeted concatemer integration events in both initial and follow-up matings. They also show that these unintended integration events can be separated from scHDR alleles in subsequent generations. However, apart from these technical peculiarities, using 5’biotinylated DNA donor fragments is a very efficient way to obtain mouse lines with targeted integration of larger (1.9 kb) DNA fragments into the mouse genome. Founders #23 and #39 were both mated with C57BL/6N wild type females to generate two independent Akr1b3^+/fx^ mouse lines (Akr1b3^+/fx^ Line #1 and Akr1b3^+/fx^ Line #2, respectively). Southern blot analysis confirmed scHDR at the *Akr1b3* locus in the F3 offspring of both lines (Figure 4D, E). A probe located outside the 3’ homology arm was used to hybridize genomic DNA cleaved with NdeI and HindIII restriction enzymes. This resulted in a 4.8 kb fragment from the wild type allele and a 2.0 kb fragment that arose from an additional HindIII restriction site upon correct integration of the donor DNA fragment (Figure 4D and Figure 4E left panel). In genomic DNA digested with AflII/NdeI, fragments of 3.3 kb and 3.4 kb were detected in wild type and scHDR alleles, respectively (Figure 4D and Figure 4E right panel). Multimeric integration would result in larger fragments (3.3 kb) plus a multiple of the donor size (1.9 kb), but this was not observed in the F3 offspring of the Akr1b3^+/fx^ Line #1 or Akr1b3^+/fx^ Line #2 (Figure 4E, right panel).

In essence, in our single-step approach, we employ the 5’ biotinylation of DNA donor ends, ensuring its monomeric state upon injection. This results in efficient scHDR and stable transmission to the subsequent generation.

## DISCUSSION

Due to their high efficiency, CRISPR/Cas approaches have revolutionized genetic engineering and created a variety of new applications. For example, they have replaced conventional gene targeting strategies for introducing targeted deletions or point mutations into the mouse genome. However, the targeted insertion of longer sequences remains a challenge. Although many advanced technologies have significantly increased the knock-in efficiency of long donor fragments, little attention has been paid to the potential pitfalls of these approaches; one of these pitfalls is the multimerization of DNA donor molecules. Tandem formation of exogenous DNA copies is inevitable after donor DNA is injected into mammalian cells; this occurs due to synthesis-dependent strand annealing (SDSA) or double-strand break repair (DSBR) (2,18,19). Multimerization of exogenous DNA copies is independent of the introduction of DSB and may be facilitated when repair mechanisms associated with non-homologous end joining (NHEJ) are induced by CRISPR/Cas induced DSB (2,27). Concatemer integration of the donor DNA fragment into the targeted gene locus can then occur, thus limiting precise scHDR (20). Concatemer integrations are difficult to identify, and the genotyping PCR design used in most studies does not allow the identification of multimers. Especially when long donor fragments are used, multimerization may go undetected, even at the target site, since the PCR assay does not amplify long concatemer insertions efficiently when both primers are located outside the donor DNA fragment in the flanking 5’ and 3’ genomic sequences (“locus-locus PCR”). In addition, PCR conditions (more than 30 cycles) may lead to *in vitro* recombination events, amplifying a DNA fragment with a sequence that appears identical to the desired scHDR integration event, even though it is not present as such at the target site (21,28).

Our study explored whether 5’biotinylation of the donor ends improves scHDR efficiency compared to unmodified donor ends in the mouse genome; we recently used this approach to generate GFP-tagged alleles in medaka (21). In the current study we used a ddPCR assay to demonstrate that the multimerization of DNA donors occurs only in response to microinjection into the cells of an embryo. Donor 5’ end modification efficiently prevents multimerization in medaka and completely inhibits multimerization in mice. This method can also be used to test the efficiency and stability of the donor.

Most existing methods to generate alleles with loxP-flanked exons rely on a two-ssODN donor strategy. This process requires two linked integration events, both of which are rather rare; it is further hampered by the short homology arms (~80 nt) of the ssODNs (29). In contrast, our strategy only requires a single recombination event and uses longer homology flanks (~400-500bp), allowing for proper orientation and integration of the donor. We have also used two highly efficient sgRNAs that, when both implement DSB cutting, remove the target exon and result in a deletion of 500–600 bp. If one of the two sgRNAs is significantly less efficient than the other, the probability of a single DSB increases, and the exon/insert itself but not the intronic homology flank will be recognized for homologous recombination. This would lead to a significantly higher likelihood that only a single loxP will be inserted. Consequently, one loxP site would be missing in the resulting targeted allele (16). In our study, all analyzed founder mice with proven scHDR harbored the complete donor fragment, including both loxP sites. We also showed that embryos and founders with concatemer integrations can be easily identified using a tandem junction PCR assay; these mice can then be segregated in subsequent generations, as demonstrated with founder #23 in our study. Donor end modifications have recently been reported to increase knock-in (KI) efficiency in immortalized mammalian cells. A study published in 2020 compares the KI efficiency of numerous dsDNA donor end modifications, including 5’biotinylation, in human cells. That study finds that a 5’C6-PEG10 modification was the most efficient and increased gene KI rates by up to five times in human cell cultures. Donor 5’biotinylation also increased KI efficiency, but not to the same extent as the 5’C6-PEG10 modification (30). Furthermore, concatemer integration at the target site was observed in a maximum of two out of 11 (18%) cell clones with the 5’C6-PEG10 modification. However, that study does not compare the concatemer integration rates of modified and unmodified donor for the 5’C6-PEG10 modification or 5’biotinylation.

Another recent study finds that 5’biotinylation of donor DNA (BIO-PCR donor) increases targeted integration when used in combination with a Cas9-Streptavidin fusion protein (Cas9-mSA). The authors explain that the Biotin-Streptavidin interaction between the BIO-PCR donor and Cas9-mSA guides the donor molecule to close proximity to the DSB site, increasing HDR efficiency by up to four times in mouse embryos compared to an unmodified PCR donor and standard Cas9 (14). That study did not explore whether this increased efficiency is due to donor recruitment to the Cas9-mSA alone or whether it could also be achieved by preventing donor multimer formation of^i^ 5’biotinylated DNA donor molecules. Our findings show that modifying the donor DNA ends alone significantly increases scHDR, even when used with a Cas9 that lacks Streptavidin coupling.

DSBs introduced by Cas9 nuclease initiate competing DNA repair pathways. However, the HDR mechanism is essential to precise integration, and the unwanted, error-prone NHEJ DNA repair mechanism must be circumvented. Notably, the HDR mechanism is only active during the late S and G2 phase; furthermore, it is suppressed by the upregulated NHEJ pathway (24). Therefore, more and more studies are exploring the possibilities of manipulating cellular DNA repair processes to enhance precise genome editing in mice, either by actively blocking NHEJ and/or actively favoring HDR (31). One promising target molecule in mammalian species is DNA ligase IV, which was inhibited by Scr7 to promote HDR at the expense of NHEJ (32). However, Scr7 had minimal effects in a later study that found RS-1 to be more superior in rabbit embryos (33). RS-1 is a HDR enhancer that stimulates the building of RAD51-DNA nucleoprotein filaments that recruit homology-directed repair templates. RAD51-enhanced interhomolog repair has also been proposed to increase zygotic genome editing in mice (34). In addition, M3814 provides pharmacological inhibition of DNA protein kinases and increases HDR efficiency in human and mouse cells (35,36). So far, there is no universal strategy to modulate DNA DSB repair mechanisms for enhancing HDR efficiency; potential modulation approaches must be evaluated for different cell types and species (31). However, it needs to be noted that analysis of concatemer integration was not addressed in any of these studies.

Taken together, our study describes an advanced donor technology that significantly increases scHDR efficiency without systemically interfering with endogenous DNA repair mechanisms, thus paving the way to clinical applications.

## MATERIALS AND METHODS

### Mouse strains

Wild type C57BL/6N mice were purchased from Charles River Laboratories (Wilmington, MA, USA). Experimental procedures were approved by the regional council Karlsruhe, Germany according to the Animal Welfare Act (AZ35-9185.81/G319/14). Microinjections were done into the cytosol of C57BL/6N two-cell stage embryos. F1 founders as well as wild type and *Akr1b3^flox/+^* offspring harboring the *Akr1b3^flox^* allele were housed in the Interfaculty Biomedical Faculty (IBF) of Heidelberg University. Mice were kept under specified pathogen-free conditions on a 12-hour light/12-hour dark cycle with water and standard food (Rod18, LASvendi GmbH, Germany) available ad libitum.

### Fish husbandry

Medaka (*Oryzias latipes*) Cab strain used in this study was kept as closed stock in accordance to Tierschutzgesetz §11, Abs. 1, Nr. 1 and with European Union animal welfare guidelines. Fish were maintained in a constant recirculating system at 28°C on a 14 h light/10 h dark cycle (Zucht- und Haltungserlaubnis AZ35-9185.64/BH).

### Cloning of donor plasmids

*Akr1b3* donor plasmid containing 5’ and 3’ homology arms, loxP sites flanking exon 3 and the additional diagnostic HindIII restriction site was generated by GoldenGATE assembly published by Kirchmaier et. al 2013 (22). Homology arms and exon 3 of the murine *Akr1b3* locus were amplified with Q5 polymerase (New England Biolabs) using genomic DNA from mouse R1 ES cells (23), primer pairs are listed in Supplementary Table 1. PCR products were digested with BsaI (New England Biolabs) followed by gel purification and ligation into respective entry vector. Entry vectors 1-5 were digested with BsaI (New England Biolabs) and ligated into the pGGDestSC-ATG destination vector (addgene #49322) according to the protocol published by Kirchmaier et al., 2013 (22). *Fam129b* and *Jpt2* donor plasmids served as PCR templates and were generated by blunt ligation of synthesized dsDNA fragments (IDT) into the pJet1.2/blunt cloning vector (Thermo Scientific).

### Generation of DNA donor fragments

For the *Akr1b3* donor PCR fragment universal primer pairs designed to amplify the donor sequence from destination vector pGGDestSC-ATG (addgene #49322) were described previously (21). Primers were synthesized by Sigma Aldrich with 5′biotin modification or without modification for control primers (Supplementary Table 1). DNA donor fragments were amplified using Q5 polymerase (New England Biolabs), PCR products (donor-5’biotin, donor-unmodified) were gel purified using the Monarch DNA Gel Extraction Kit (New England Biolabs, T1020S) and eluted in 20μl embryo transfer water (Sigma, W1503) prior to microinjection.

### sgRNA design

sgRNAs were designed with the CRISPR/Cas9 target online predictor tool CCTop with standard parameters (PAM = NGG, target site length = 20 nt, core length = 12 nt, max mismatches = 4, mouse genome = GRCm38/mm10) (24). The specific target sites were selected to induce DSBs in the intronic 5′ and 3′ regions of the target exon, respectively. The target sites on the donor were replaced by loxP sites, which prevented cutting of the dsDNA donor. Generation of sgRNA constructs and *in vitro* transcription was carried out according to the protocol of Stemmer et al., 2015 (24). Purchased sgRNA oligos were annealed and cloned into vector DR274 (25) (DR274 was a gift from Keith Joung, Addgene plasmid #42250) via the BsaI restriction site. The resulting sgRNA plasmid was linearized by FastDigest DraI (Thermo Scientific) and served as template for sgRNA *in vitro* transcription using the T7 MEGAshortscript Kit (Thermo Scientific, AM1354) according to the manufacturer’s protocol. sgRNAs were purified using the RNeasy Kit (Qiagen, 74104). For quality control 450 ng of sgRNA was mixed with 2x RNA loading buffer and loaded onto a 1.5 % TAE gel. sgRNA integrity was confirmed by a band at 200 bp of the used DNA marker.

### Generation of *Cas9* mRNA

*heiCas9* mRNA (26) was generated by *in vitro* transcription using the mMESSAGE mMACHINE™ Sp6 transcription kit according to the manufacturer’s protocol (Thermo Scientific, AM1340). *heiCas9* mRNA was purified using the RNeasy Kit (Qiagen, 74104). mRNA quality was confirmed using a test gel. 400 ng of *heiCas9* mRNA was mixed with 2x RNA loading buffer and loaded onto a 1.5 % TAE gel. High mRNA integrity was assumed by a smear-free band on the level of the 2088 bp band of the used DNA marker.

### Microinjection of medaka zygotes

One-cell stage wild-type medaka (*Oryzias latipes*) Cab strain zygotes were microinjected into the cytoplasm with injection mixes containing 5 ng/μl unmodified or 5 ng/μl 5’biotinylated PCR donor (*Akr1b3-*flox donor fragment, see Supplementary Figure 2 diluted in nuclease free water. After injection, embryos were raised in 1× ERM (10× ERM stock: 170 mM NaCl, 4 mM KCl, 2.7 mM CaCl_2_.2H_2_O, 6.6 mM MgSO_4_.7H_2_O, 170 mM HEPES) for 1 or 6 hours prior to ddPCR analysis.

### Microinjection of mouse embryos

An injection mix according to the protocol of Gu et al., 2018 was prepared containing *heiCas9* mRNA (100 ng/μl), sgRNAs (each 50 ng/μl) and PCR donor-5′biotin or unmodified PCR donor (donor-unmod) (20 ng/μl) dissolved in injection buffer (10 mM Tris-HCl, 0.25 mM EDTA, pH 7.4) (14). For the analysis of donor multimerization in Figure 2 embryos were injected with a mix without *heiCas9* mRNA and sgRNAs. To prevent cannula clogging the injection mix was filtered shortly before the microinjection using a Corning® Costar® Spin-X®centrifuge tube-filter 0.22μm (Sigma Aldrich, CLS8160). Zygotes were collected from timed matings and cultured to the two-cell stage *in vitro* in M2 medium until and during the injection. The injection into each blastomere of the two-cell stage embryo was carried out by carefully inserting the injection needle at the equatorial level into the cytoplasm of each embryo. About 1-2 picoliters were injected. Thereafter, the two-cell stage embryos were placed in preincubated KSOM medium under paraffin oil for 1-2 hours to select the lysed and intact two-cell stage embryos. For the *in vitro* experiments, blastocysts were cultured in KSOM medium covered with paraffin oil in 5% CO_2_, 38.0°C at 98 % atmospheric humidity in the incubator. For the *in vivo* experiments, the two-cell stage embryos were transferred via oviduct transfer two hours after the injection in 0.5 days pseudo pregnant foster mothers.

### Droplet digital PCR

Droplet digital PCR (ddPCR) was used to determine multimerization of 5′biotinylated donor molecules after injection into medaka and mouse embryos. To quantify the level of multimerization the ratio between detected tandem junctions and total donor molecules was calculated. Therefore, ddPCR assays for tandem junctions and total donor molecules were designed which enabled the independent detection of multimerization based on FAM labelled probes (probe 2^a^ and probe 2^b^, Figure 2C and Supplementary Table 1) and total donor molecules based on a HEX labelled probe (probe1, Figure 2C and Supplementary Table 1). ddPCR assay probe1 detects all donor molecules independent form multimerization state. ddPCR assay probe 2^a^ + probe 2^b^ detects all possible tandem junctions independent from orientation of the donor multimer (head-to-head, head-to-tail or tail-to-tail, Supplementary Figure 7). Therefore, one FAM labelled probe was placed on the bottom strand within the 3′homology arm (probe ^2a^ in Figure 2C) of the donor and one on the top strand within the 5’ homology arm (probe ^2b^ in Figure 2C) of the donor.

DNA extraction from injected medaka (*Oryzias latipes*) embryos followed one hour and six hours after microinjection. Medaka embryos were transferred into 80 μl of 50 mM NaOH, chorions were mechanically crushed with individual pestles and embryos were incubated at 90°C for 20 min followed by the addition of 20 μl of 50 mM TrisHCl. DNA from two-cell stage mouse embryos was extracted by transferring the embryos one hour and six hours after injection into 5 μl lysis buffer (DirectPCR buffer, Viagen, supplemented with 2.5 μg/ml Proteinase K, AppliChem) incubated at 55°C for 15 min followed by 85°C for 15 min. Donor dsDNA of medaka and mouse was digested with BamHI (in loxP sites) for 1h at 37°C to ensure splitting of multimers into tandem junctions, hence enabling the distribution of tandem junctions into individual droplets. The ddPCR multiplex mix was prepared by adding 11 μl of 2x Supermix for Probes (no UTPs, BioRad), 1.1 μl of all four primers (900 nM), 1.1 μl of probe (250 nM) as well as 1.1 μl – 3.3 μl of embryo lysate. ddPCR multiplex mix was filled up with ddH_2_O to a total volume of 22 μl per reaction.

Droplets were generated using a QX200 Droplet Generator (Bio Rad) following the manufacturer’s protocol. After transferring the generated droplets to a 96-well plate (Thermo Scientific) an end point PCR was performed using the C1000 Touch PCR thermal cycler (Bio Rad) using the following protocol: 95°C for 10 min, 30 cycles of 95°C for 30 s, 54°C for 20 s and 72°C for 2 min. Droplets were read out for FAM and HEX positive events using the QX200 droplet reader (Bio-Rad) followed by analyzing data with the QuantaSoft™ program (Bio-Rad, version 1.7.4.0917). The concatemer index was defined as tandem junction event counts/donor molecule event counts.

### Genotyping

In the first series of experiments, the injected two-cell stage embryos were cultured *in vitro* for 3.5 additional days until E4.5 blastocyst stage. Pools of 2 or 3 E4.5 blastocysts were lysed in 10 μl lysis buffer (DirectPCR buffer, Viagen, supplemented with 2.5 μg/ml Proteinase K, AppliChem) and incubated at 55°C for 15 min followed by 85°C for 15 min. In a second approach, the injected two-cell stage embryos were transferred via oviduct transfer to allow implantation into foster mothers, and E14.5 embryos were isolated and entirely lysed in 60 μl lysis buffer and incubated at 55°C for 2 hours followed by 85°C for 15 min. In a third set of experiments, foster mothers that underwent oviduct transfer were allowed to deliver, and ear biopsies were taken in the adult offspring, lysed in 100 μl lysis buffer and incubated at 55°C overnight followed by 85°C for 45 min.

Genotyping was performed by Q5 PCR (0.5 μl Q5® HiFi DNA Polymerase (New England Biolabs, M0491), 1xQ5 reaction buffer, 200 μM dNTPs, 500 nM primer forward and reverse and 0.5-1.0 μl genomic DNA) with a locus-locus primer pair located outside the 5′ and 3′ homology arms of the integrated donor DNA fragment and 5′and 3′junction primer pairs (Supplementary Table 1). The following Q5 HiFi PCR protocol was applied to identify scHDR, targeted integration and potential concatemer integration at different loci: 98°C 30 sec denaturation, 30 cycles of 98°C 10 sec denaturation, annealing at 60-69°C for 30 sec and elongation at 72°C for 60-120 sec. PCR products were mixed with 5x DNA loading dye (Thermo Scientific, R0611) and separated on a 1% agarose gel.

### Concatemer detection

Tandem PCR was carried out to detect donor concatemer integrations in the mouse genome. Therefore, the forward primer (F2 in Figure 1A) was placed in the 3′loxP site of the donor whereas the reverse primer (R2 in Figure 1a) was located in the 5′loxP sequence of the donor resulting in an inverse primer orientation (Supplementary Table 1). Tandem PCR was performed by Q5 PCR (0.5 μl Q5® HiFi DNA Polymerase (New England Biolabs, M0491), 1x Q5 reaction buffer, 200 μM dNTPs, 500 nM primer forward and reverse and 0.5-1.0 μl genomic DNA). The following PCR protocol was applied to identify concatemer integrations in the mouse genome: 98°C 30 sec denaturation, 30 cycles of 98°C 10 sec denaturation, annealing at 65-67°C for 30 sec and elongation at 72°C 60 sec. PCR products were mixed with 5x DNA loading dye (Thermo Scientific, R0611) and separated on a 1% agarose gel.

### Southern Blot

Genomic DNA was isolated from the liver of Akr1b3^+/fx^ (F3 generation offspring) and Akr1b3^+/+^ littermates using the phenol-chloroform extraction method. The Southern blot was carried out by Celplor LLC (Cary, NC USA). Aliquots of 10 μg genomic DNA were digested with HindIII/NdeI and AfIII/NdeI restriction enzymes, respectively, at 37°C for approximately 24 hours with 10 units of each enzyme. After digestion, an aliquot of each digest was visualized on an agarose gel to ensure genomic DNAs were completely digested. After transfer of the DNA the blots were probed with a DNA fragment (sequence see Supplementary Figure 8) located outside of the donor DNA in the intron downstream of exon 4 and 5’ of the diagnostic NdeI restriction site. Experimental procedures including non-radioactive probe generation and labelling, Southern blotting and signal recording are performed following SOPs of Celplor LLC.

### Sequencing

PCR products were sequenced by the service company Eurofins Genomics (Ebersberg, Germany) using primers F1 and R1 listed in Supplementary Table 1.

### Statistics

Statistical analyses were performed using OriginPro 2019 and Excel 2010. Data are expressed as mean ± standard deviation (SD). Normality distribution of the data was tested using a Shapiro-Wilk test and considered normally distributed if p > 0.05. Difference between 2 groups was tested using a two-sided t-test. P values < 0.05 were considered statistically significant.

## AVAILABILITY

All data that support the findings of this study are available in this study within the manuscript and/or its supplementary materials.

## SUPPLEMENTARY DATA

Additional material and data can be found in the Supplementary Information.

## ACKNOWLEDGEMENT

We are thankful to Christin Richter, Hans-Peter Gensheimer, Beate Hilbert and the entire team from the Interfakultäre Biomedizinische Forschungseinrichtung (IBF) from Heidelberg University for expert technical assistance. We thank Keith Joung for the DR274 plasmid.

## AUTHOR CONTRIBUTIONS

Rebekka Medert, Conceptualization, Data curation, Formal analysis, Validation, Investigation, Visualization, Writing—original draft, Writing—review and editing

Thomas Thumberger, Conceptualization, Formal analysis, Supervision, Validation, Investigation, Methodology, Writing—review and editing

Tinatini Tavhelidse, Conceptualization, Formal analysis, Supervision, Validation, Investigation, Visualization, Writing—review and editing

Tobias Hub, Methodology, Investigation, Data curation

Yoko Oguchi, Investigation, Data curation

Tanja Kellner, Methodology

Frank Zimmermann, Methodology; Investigation; Validation, Formal analysis

Sascha Dlugosz, Methodology; Investigation; Validation, Formal analysis

Joachim Wittbrodt, Conceptualization, Formal analysis, Supervision, Funding acquisition, Investigation, Writing—review and editing

Marc Freichel, Conceptualization, Formal analysis, Supervision, Funding acquisition, Investigation, Writing-original draft, Project administration, Writing—review and editing

## Supplementary Information

**Supplementary Table 1.**
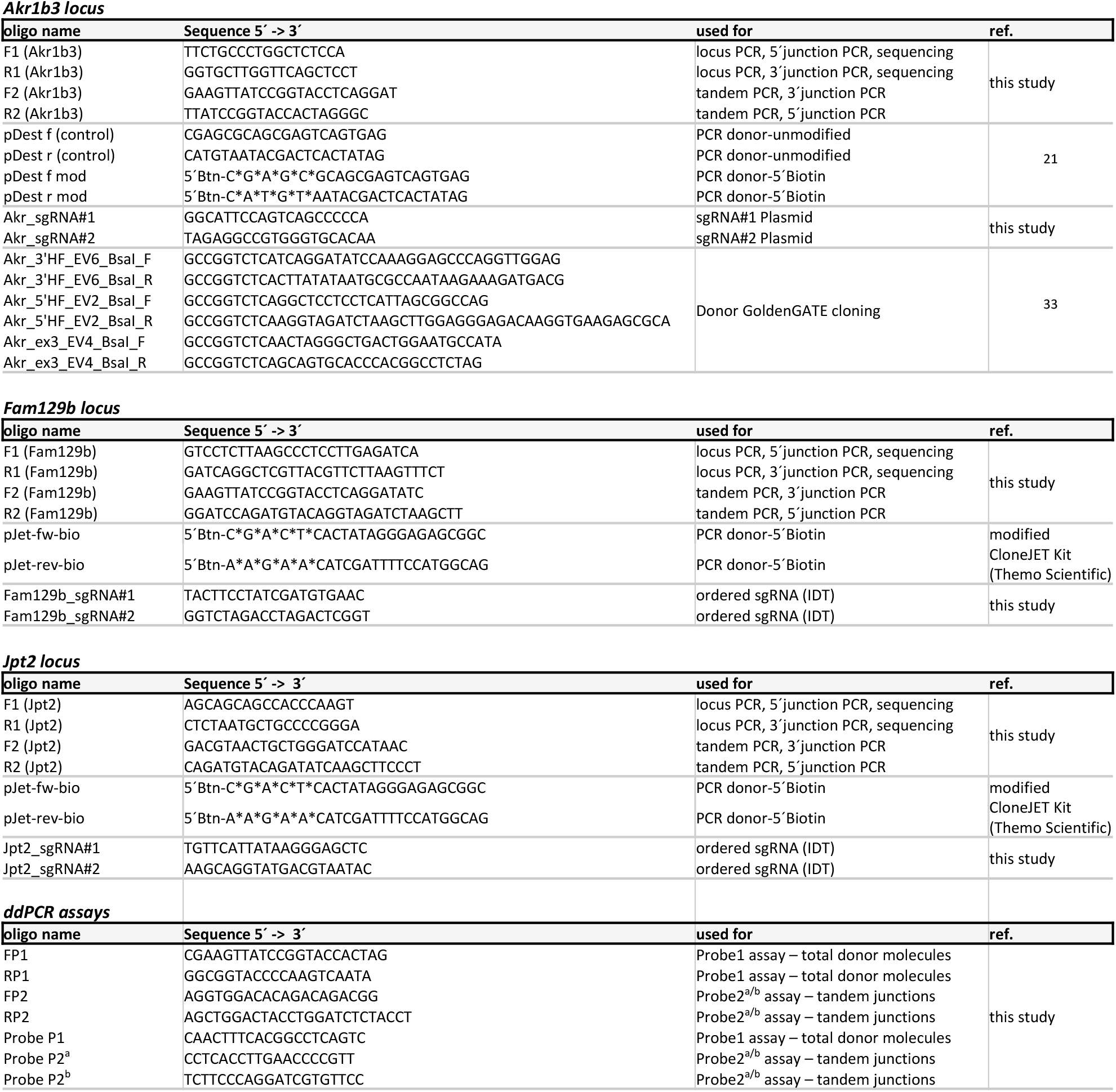
List of oligonucleotide sequences used in this study. Asterisks indicate phosphorothioate bonds.

**Supplementary Figure 1.**
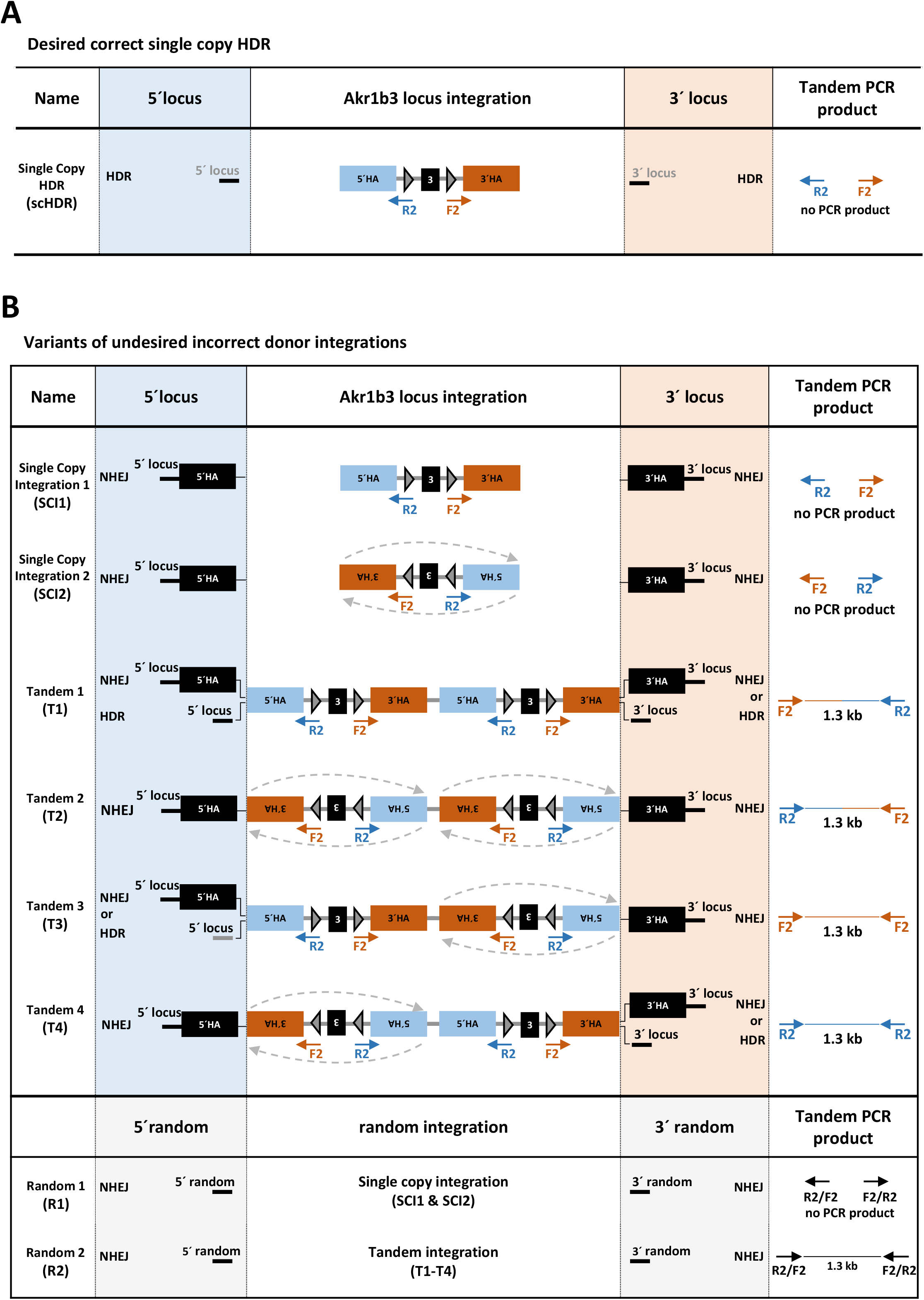
Possible precise and imprecise integration events of the dsDNA donor at the *Akr1b3* locus. Tandem PCR strategy was used to identify the integration of donor concatemers in the mouse genome. The forward primer (F2; orange arrow) is located at the 3′end of the donor, whereas the reverse primer (R2; blue arrow) is located at the 5′end. **(A)** Correct scHDR results in no tandem PCR product. **(B)** Imprecise single copy integration results also in no tandem PCR product (SCI1 & SCI2), whereas tandem variants T1-T4 and random integration variant R2 generate a tandem PCR product of about 1.3 kb.

**Supplementary Figure 2.**
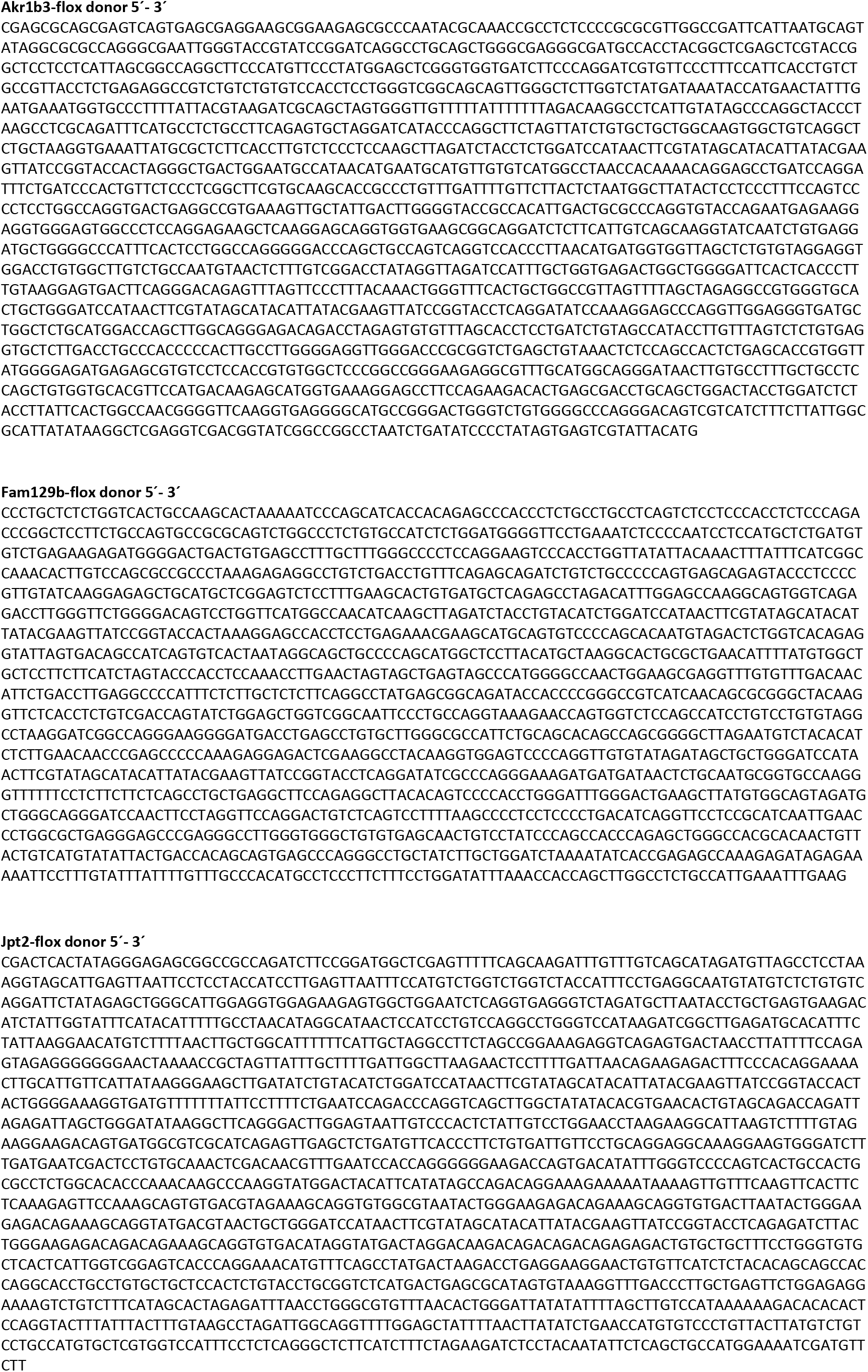
Donor sequences used in this study.

**Supplementary Figure 3.**
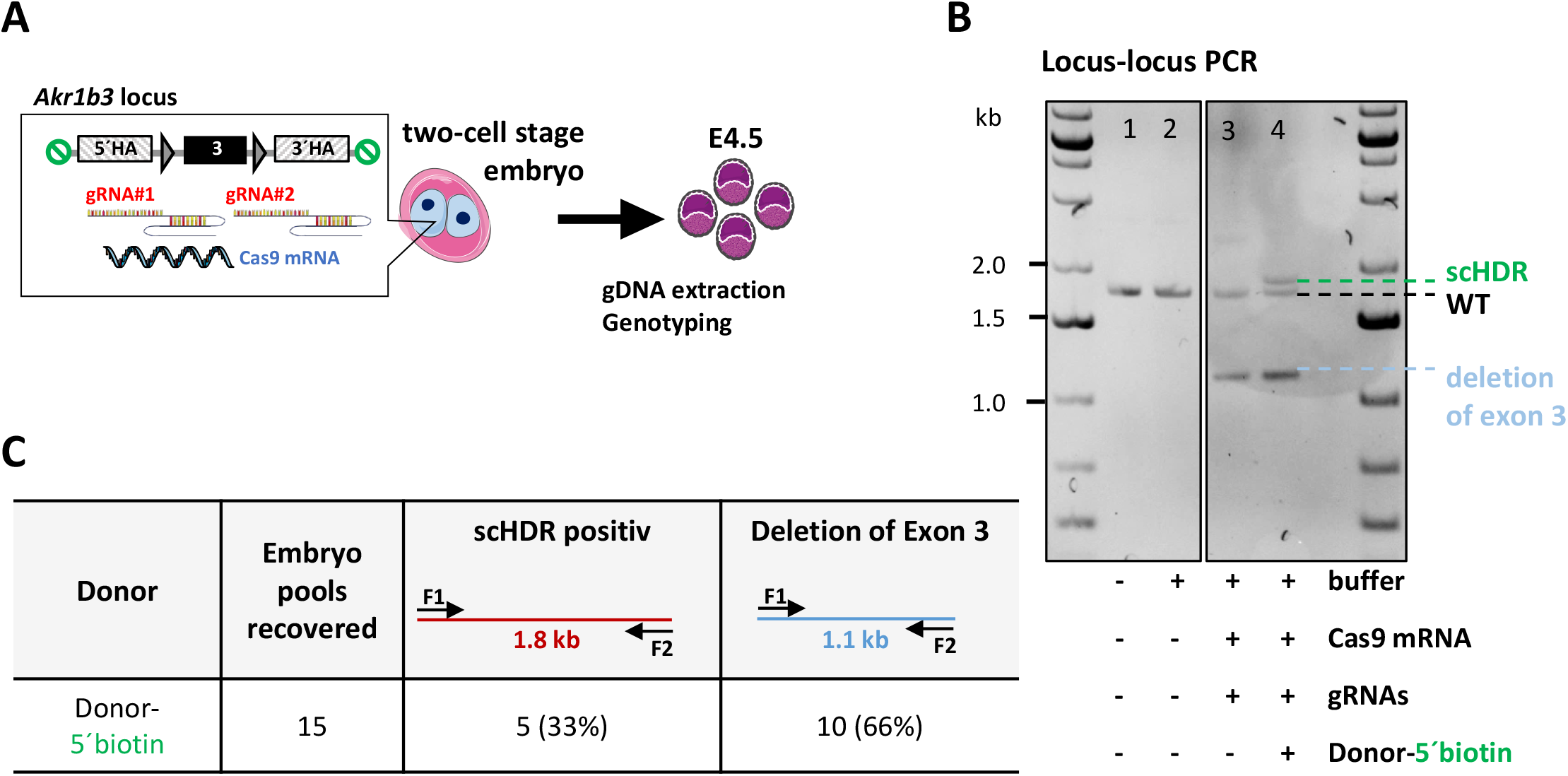
Genotyping of E4.5 embryos demonstrate scHDR of a 5′biotinylated dsDNA donor. **(A)** Two-cell stage microinjection strategy to deliver *Cas9* mRNA, sgRNAs and PCR donor-5’biotin into the mouse embryo followed by genotyping at embryonic stage E4.5. Two to three blastocysts were pooled on 5 independent microinjection days. **(B)** Locus-locus PCR genotyping performed in pools of 2-3 E4.5 embryos confirms high efficiency of both sgRNAs resulting in the deletion of exon 3 (lanes 3 and 4) and identifies CRISPR/Cas9 mediated scHDR in the presence of PCR donor-5’biotin (lane 4). **(C)** Efficacy of single copy integration analyzed at blastocyst stage E4.5.

**Supplementary Figure 4.**
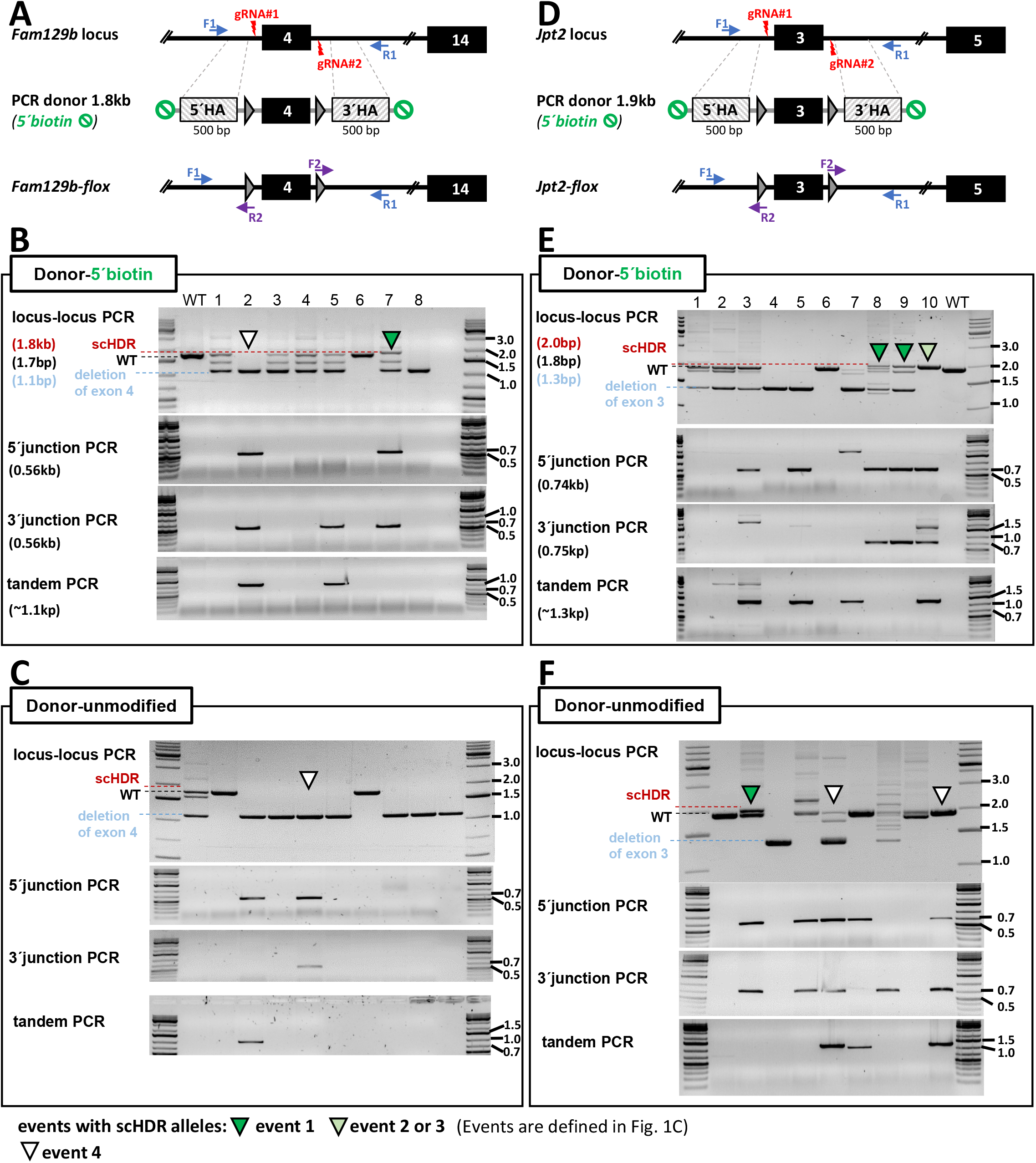
Donor 5′biotin mediates efficient scHDR at the *Fam129b* and *Jpt2* locus. **(A-C)** Targeting strategy and genotyping of the *Fam129b* locus. **(A)** Two-sgRNA targeting strategy for the generation of a scHDR allele using a dsDNA donor template containing exon 4 flanked by loxP sites. **(B)** CRISPR/Cas9 mediated scHDR at the *Fam129b* locus is only present when PCR donor-5‘biotin is injected. **(C)** Genotyping PCRs identified tandem junctions but no scHDR events upon injection of PCR donor-unmodified. **(D-F)** Targeting strategy and genotyping of the *Jpt2* locus. **(D)** Two-sgRNA targeting strategy for the generation of a scHDR allele using a dsDNA donor template containing exon 3 flanked by loxP sites. **(E)** CRISPR/Cas9 mediated scHDR at the *Jpt2* locus was increased when PCR donor-5‘biotin was injected. **(F)** Genotyping PCRs identified tandem junctions and scHDR events in the presence of PCR donor-unmodified. scHDR efficiency was evaluated at embryonic stage E14.5. Quantification of scHDR efficiency at the *Fam129* and *Jpt2* locus is displayed in Figure 3E.

**Supplementary Figure 5.**
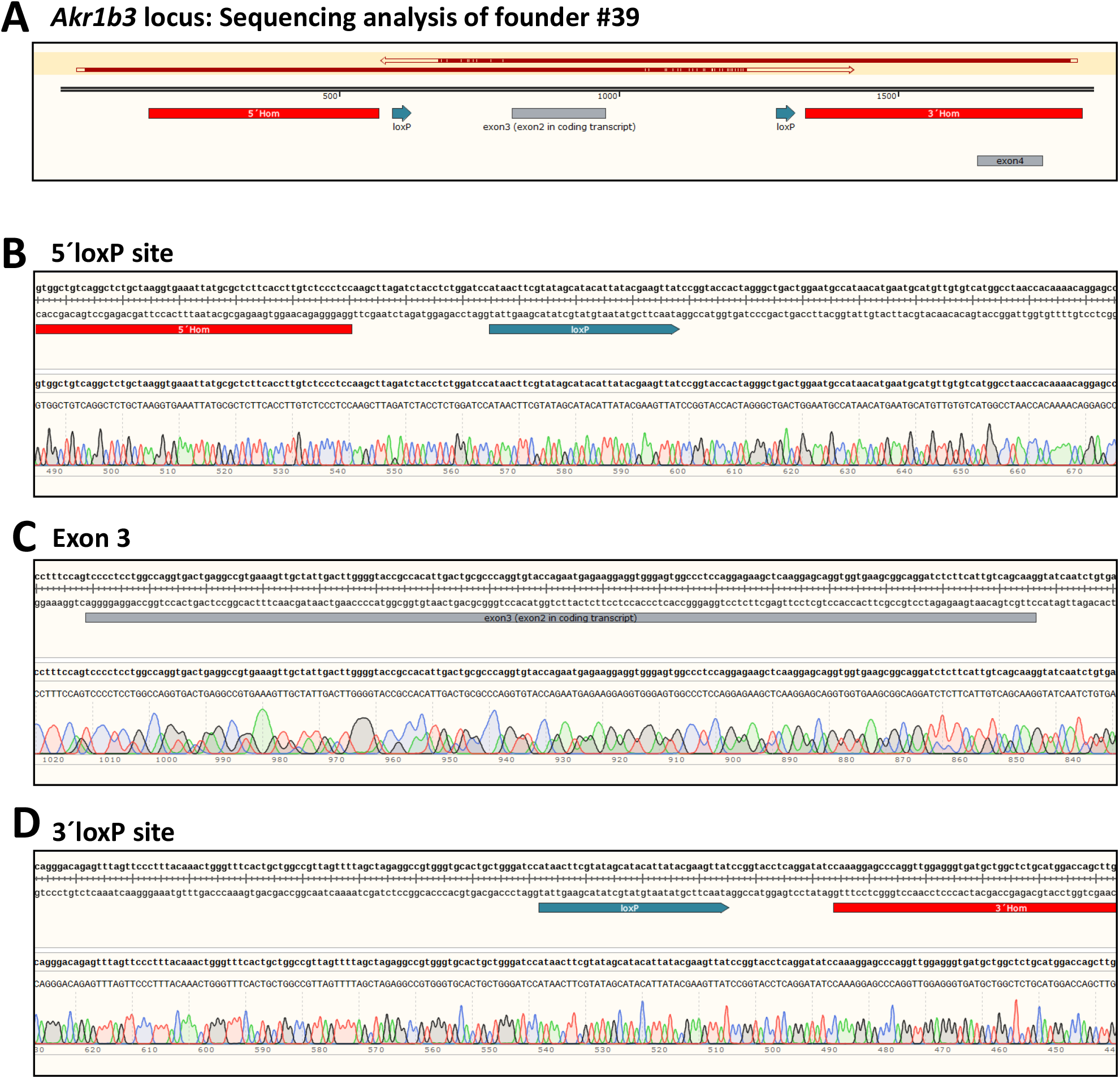
Sequencing of scHDR locus PCR product of *Akr1b3-flox* founder #39. **(A)** Selection of the *Akr1b3* locus containing exon 3 is shown together with the homology arms and location of loxP sites. **(B-D)** Locus-locus PCR product obtained by primers F1/R1 and genomic DNA of founder #39 as template was sequenced. DNA sequence around 5‘loxP site **(B)** covering exon 3 and adjacent introns **c**, and around the 3‘loxP site **(D)** showed correct integration (scHDR) in founder #39.

**Supplementary Figure 6.**
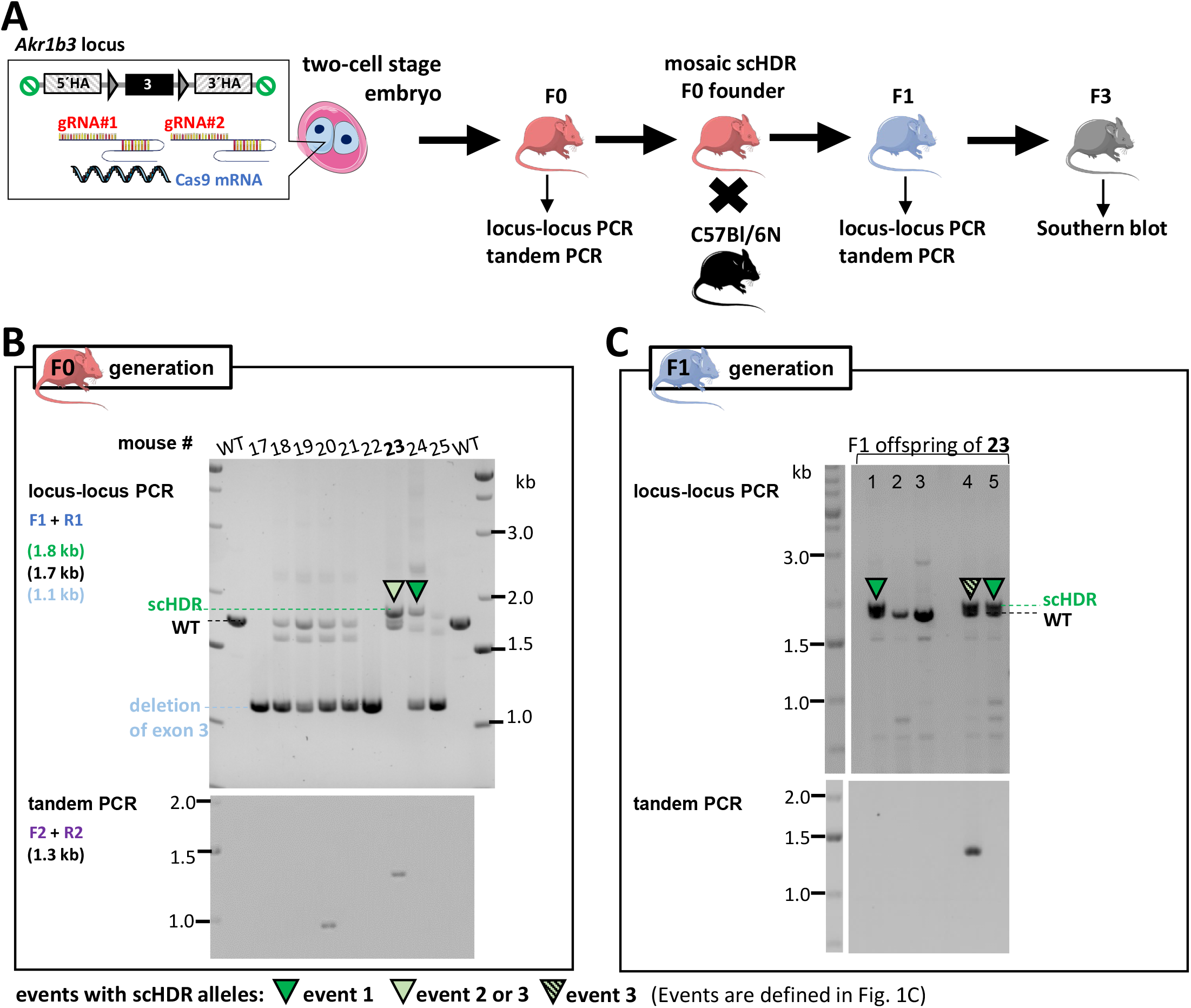
F0 founder positive for tandem junction PCR can still exhibit and inherit an allele with precise genome editing (scHDR). **(A)** Two-cell stage microinjection strategy to deliver *Cas9* mRNA, sgRNAs and PCR donor-5‘biotin into the mouse embryo followed by genotyping of single founders followed using ear punch biopsies. Mating strategy of mosaic scHDR positive founder and genotyping of F1 generation. Southern Blot analysis in F3 generation confirms scHDR events. **(B)** Locus-locus PCR (assay #1) of F0 mice identified CRISPR/Cas9 mediated scHDR alleles (dark and light green arrowheads) upon injection of a PCR donor-5‘biotin. Tandem junction PCR of F0 mice identified concatemer integration of the PCR donor-5‘biotin. Concatemers are also detected in the presence of a scHDR allele in founder #23 (light green arrowhead). **(C)** Locus-locus PCR of F1 generation of founder #23 identified germline transmission of scHDR alleles (dark and light green with back stripes arrowheads). Tandem junction PCR uncovered off-target concatemer integration of the donor in scHDR positive F1 offspring (light green with black stripes arrowhead).

**Supplementary Figure 7.**
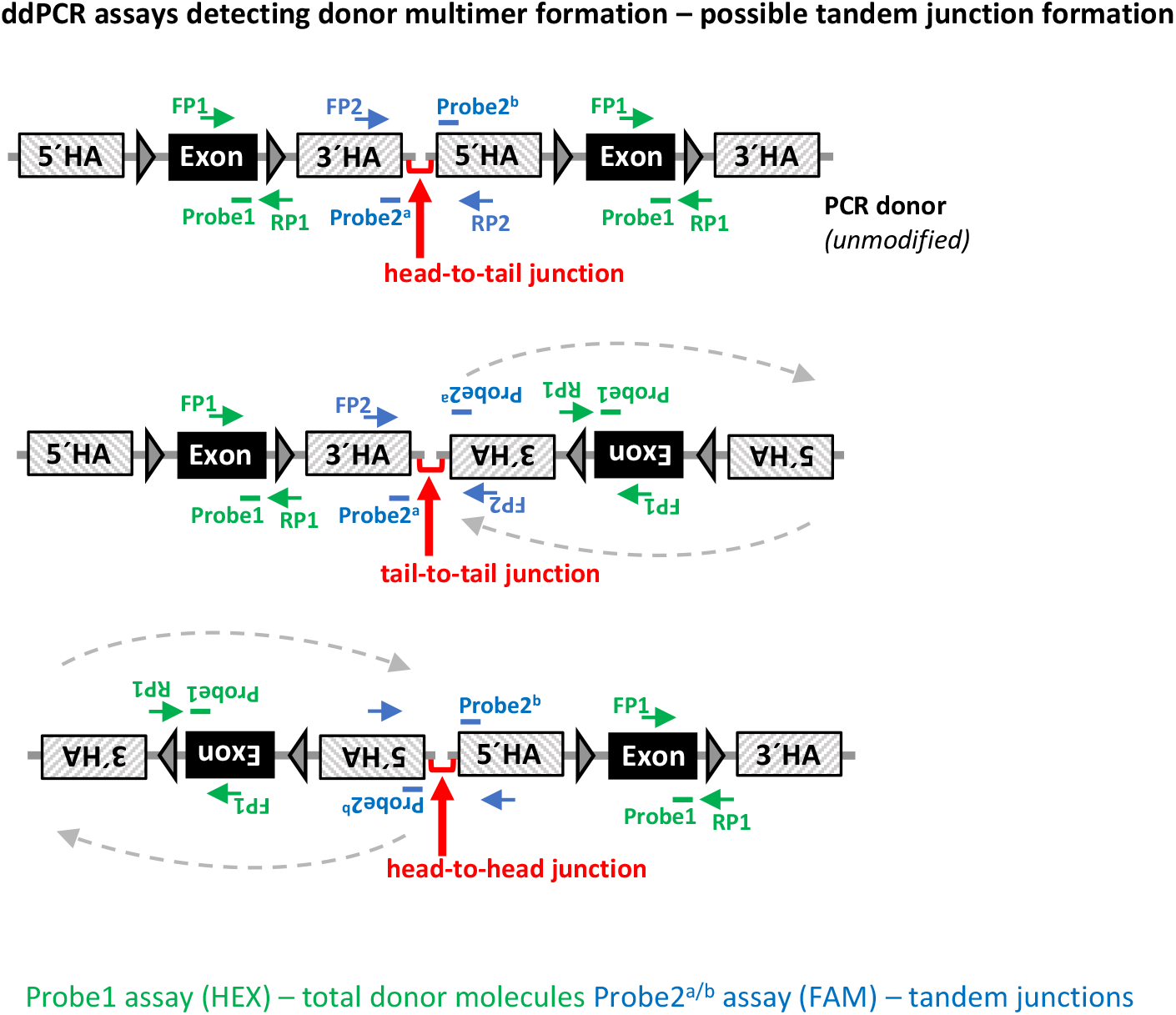
Possible multimerization junctions of donor molecules in medaka and mouse embryos. Donor molecules can form head-to-tail (upper variant), tail-to-tail (middle variant) and head-to-head (lower variant) tandem junctions. Droplet digital PCR (ddPCR) assays to quantify total donor molecule (Probe 1 assay, HEX) and tandem junctions (Probe 2 assay, FAM) resulting from head-to-tail, tail-to-tail and head-to-head donor multimerization.

**Supplementary Figure 8.**
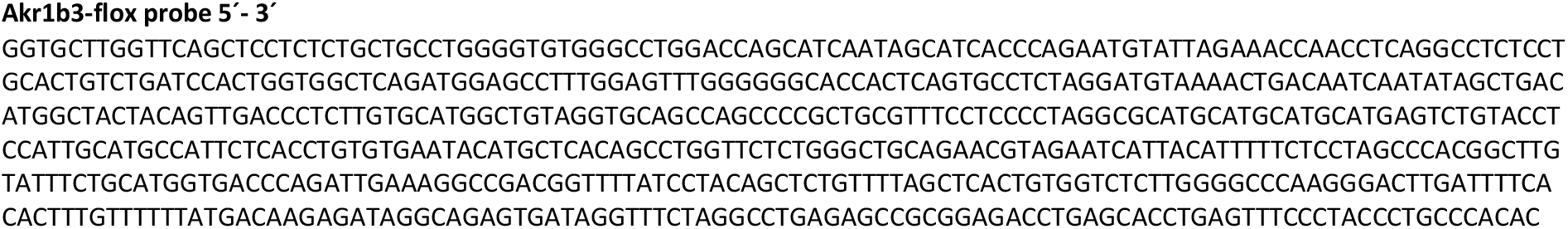
Sequence of *Akr1b3-flox* probe used for Southern blot analysis.

